# BioCantor: a Python library for genomic feature arithmetic in arbitrarily related coordinate systems

**DOI:** 10.1101/2021.07.09.451743

**Authors:** Pamela H. Russell, Ian T. Fiddes

## Abstract

**Motivation:** Bioinformaticians frequently navigate among a diverse set of coordinate systems: for example, converting between genomic, transcript, and protein coordinates. The abstraction of coordinate systems and feature arithmetic allows genomic workflows to be expressed more elegantly and succinctly. However, no publicly available software library offers fully featured interoperable support for multiple coordinate systems. As such, bioinformatics programmers must either implement custom solutions, or make do with existing utilities, which may lack the full functionality they require.

**Results:** We present BioCantor, a Python library that provides integrated library support for arbitrarily related coordinate systems and rich operations on genomic features, with I/O support for a variety of file formats.

**Availability and implementation:** BioCantor is implemented as a Python 3 library with a minimal set of external dependencies. The library is freely available under the MIT license at https://github.com/InscriptaLabs/BioCantor and on the Python Package Index at https://pypi.org/project/BioCantor/. BioCantor has extensive documentation and vignettes available on ReadTheDocs at https://biocantor.readthedocs.io/en/latest/.

## Introduction

The term “genomic feature arithmetic” refers to coordinate operations on representations of genomic features such as genes, transcripts or non-coding elements. Examples of feature arithmetic operations include coordinate conversion between coordinate systems, binary set theoretic operations such as intersection or union of features, and unary operations such as iterating over windows of a feature or reversing the strand of a feature. A variety of computational tools exist that support feature arithmetic operations, both as command line utilities (Bedtools^1^; BEDOPS ^2^) and software libraries (Pybedtools ^3^; PyRanges ^4^; the GenomicFeatures^5^ BioConductor package). However, no library exists that supports rich feature arithmetic operations across arbitrarily related coordinate systems: for example, a series of nested coordinate systems including exon-relative, transcript-relative, and chromosome-relative coordinates.

Here we present BioCantor, a Python library implementing a rich set of feature arithmetic operations including many not supported by other packages (**Table 1**). BioCantor facilitates importing and exporting common annotation file formats into a simple genome annotation data model, supports arbitrarily related coordinate systems and abstracts coordinate conversion from the user.

**Table 1.**
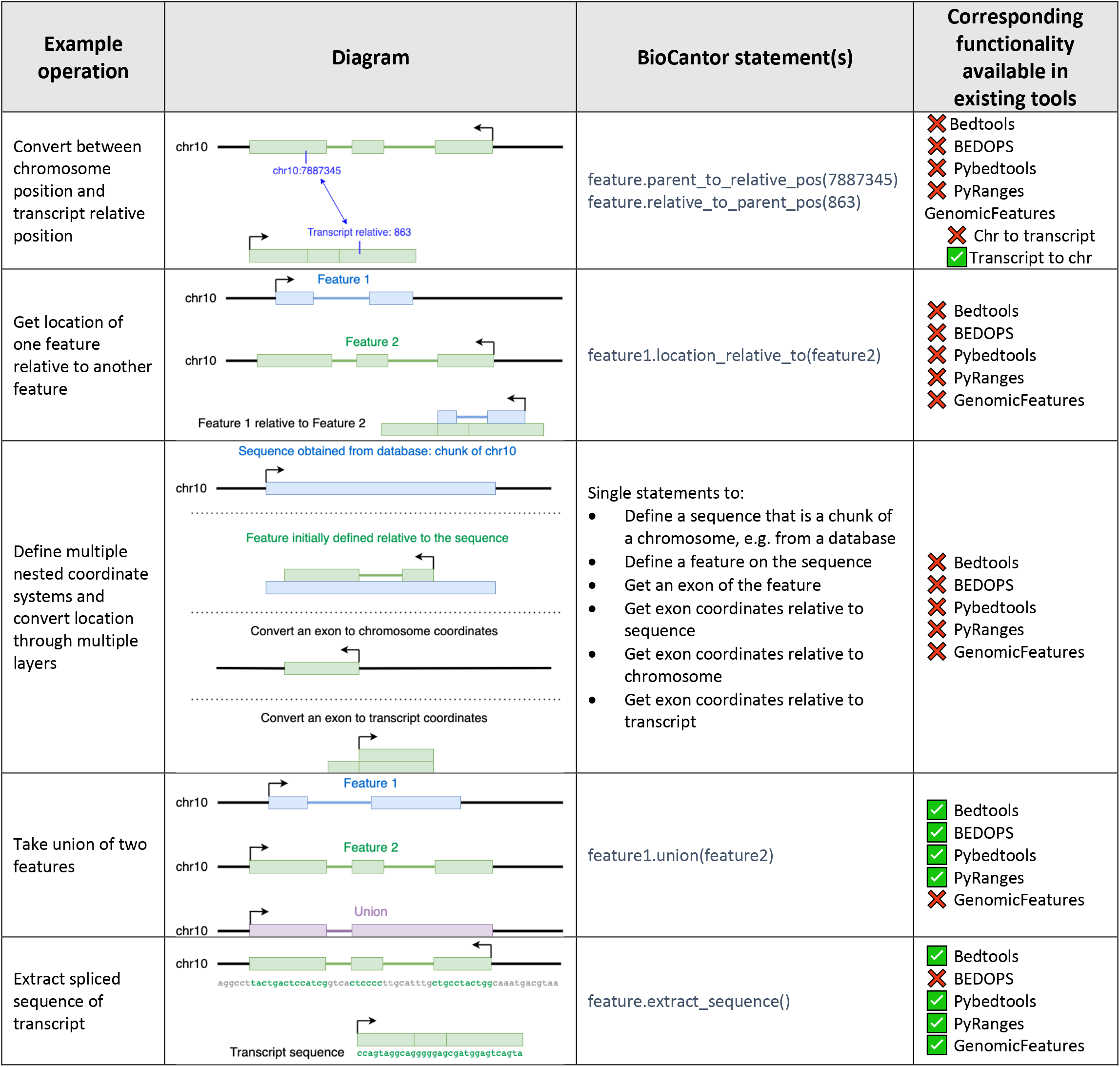
Example BioCantor operations and availability in existing tools.

## Results

### BioCantor paradigm

The basis of the BioCantor paradigm is that objects are linked by parent/child relationships. Once a parent/child hierarchy is established, coordinate operations can move around the hierarchy with this detail abstracted to the user; for example, converting a feature annotation from one reference sequence to another. In most cases, parent/child relationships are used to establish the parent as the frame of reference for the location of the child, though these relationships may exist without defining a coordinate system.

Three main object types populate this paradigm. `Location` objects represent blocked and stranded features. `Sequence` objects hold sequence data. `Parent` objects define parent/child relationships. `Sequence` and `Parent` are concrete classes. `Location` is an abstract class with three implementations: the singleton `EmptyLocation`, `SingleInterval` (a contiguous interval with start and end coordinates), and `CompoundInterval` (a multi-block feature).

All objects hold pointers to their own optional parent. `Parent` objects do not hold pointers to children and can be reused for multiple children. `Location` objects, `Sequence` objects, and `Parent` objects can all have parents. Multilevel hierarchies are established when `Parent` objects have their own parents.

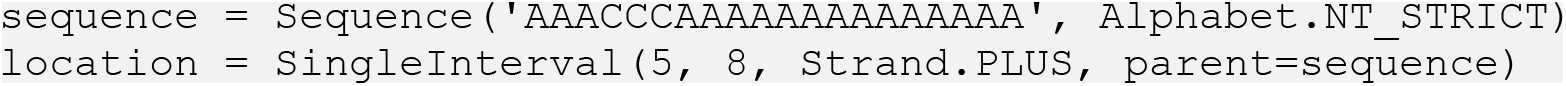
Example: instantiating a Location object that refers to a parent sequence.

The `Parent` class is very flexible in order to accommodate different types of relationships. For example, a `Parent` object can optionally hold a pointer to a `Sequence`, meaning that sequence is the frame of reference for an object with that `Parent`. A `Parent` object can optionally hold a `Location`, meaning that is the location of the child relative to that parent. `Parent` has several optional parameters which enable different types of relationships and operations.

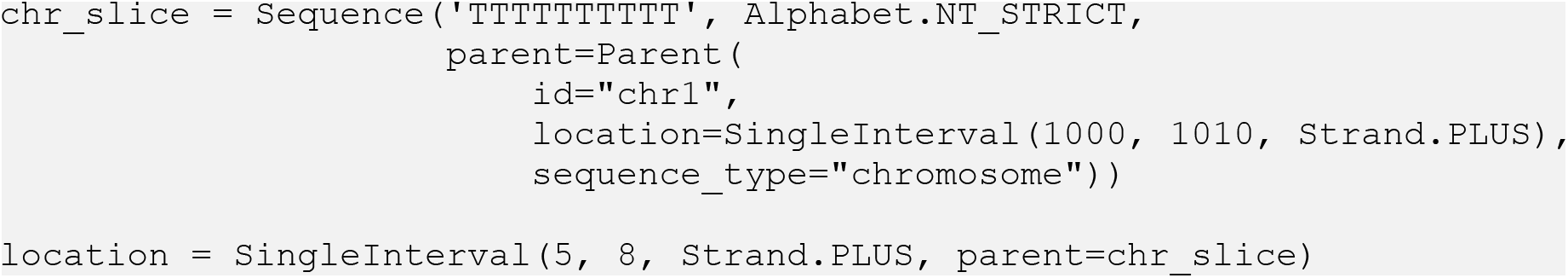
Example: a Location points to a slice of a chromosome as its parent. The chromosome slice holds sequence data. Additionally, the chromosome slice has its own Parent representing the location of the slice relative to a chromosome.

### Coordinates and coordinate conversion

`Location` classes represent blocked, stranded features with block coordinates represented by zero-based, end exclusive coordinates. These classes provide a variety of conversion methods: coordinates and features can be converted between any coordinate systems in the hierarchy with a single statement. In particular, seamless conversion between genomic and transcript-relative coordinates enables expressive statements operating in transcript space.

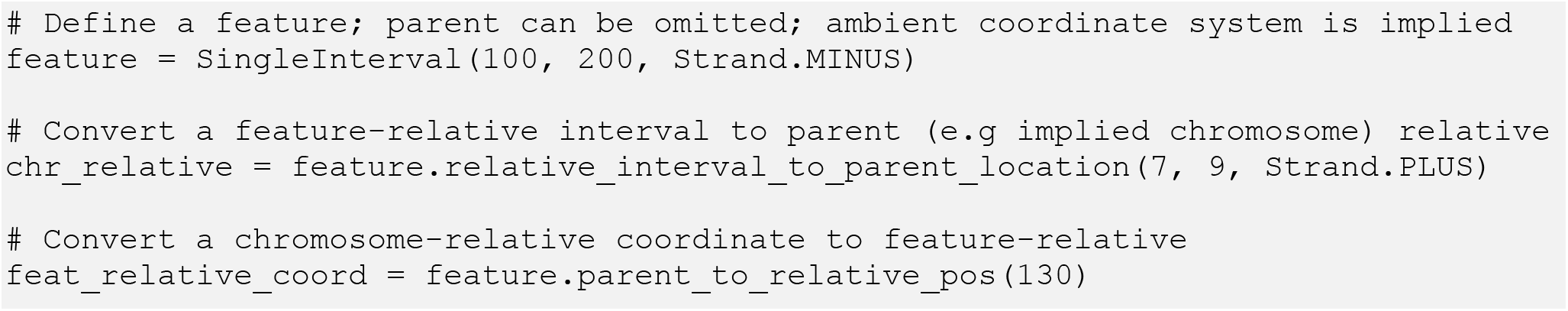
Example: converting between transcript-relative and chromosome-relative coordinates.

### Feature arithmetic

Alongside the support for coordinate systems, feature arithmetic functionality includes standard set theoretic operations (intersection, union, contains, overlaps, etc.) and other useful location operations. Splicing and strand are handled seamlessly. Moreover, the library includes special support for transcripts, coding sequences, codons, and translation, allowing users to quickly navigate among these features and retrieve their sequences.

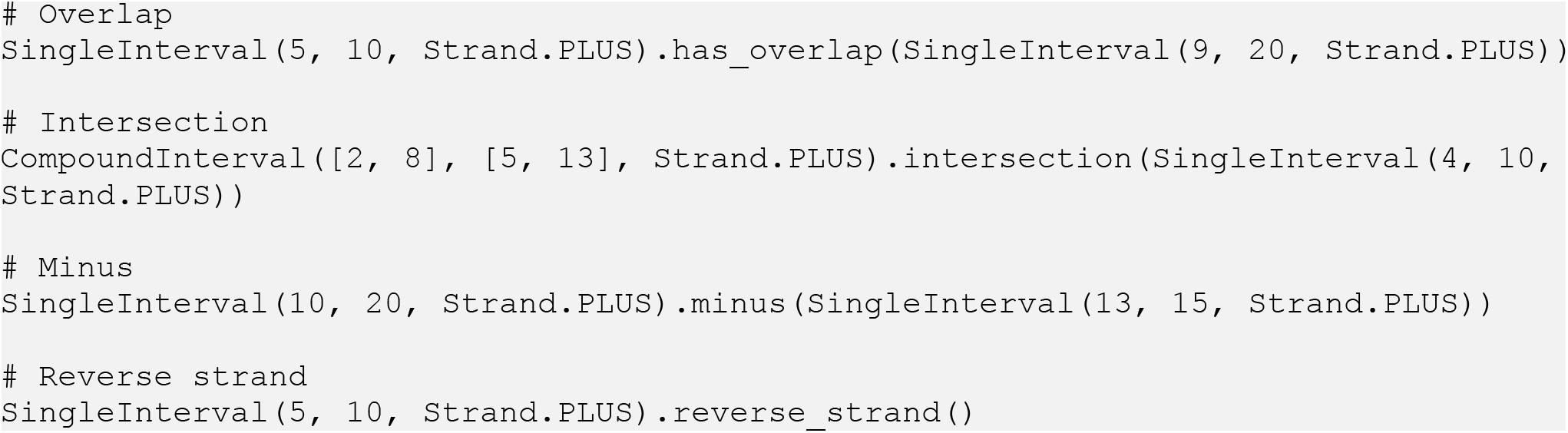
Example: feature arithmetic operations.

### Feature collections

BioCantor provides container classes to combine sets of transcripts into a gene (`GeneInterval`), sets of arbitrary features into a collection, (`FeatureIntervalCollection`), and sets of genes and/or features into an arbitrary collection (`AnnotationCollection`) **(Figure 1)**. For example, an `AnnotationCollection` object could represent the full annotation of a chromosome loaded in from a GenBank or GFF3 file. `AnnotationCollection` objects can be queried for subsets of features overlapping specific coordinates.

**Figure 1.**
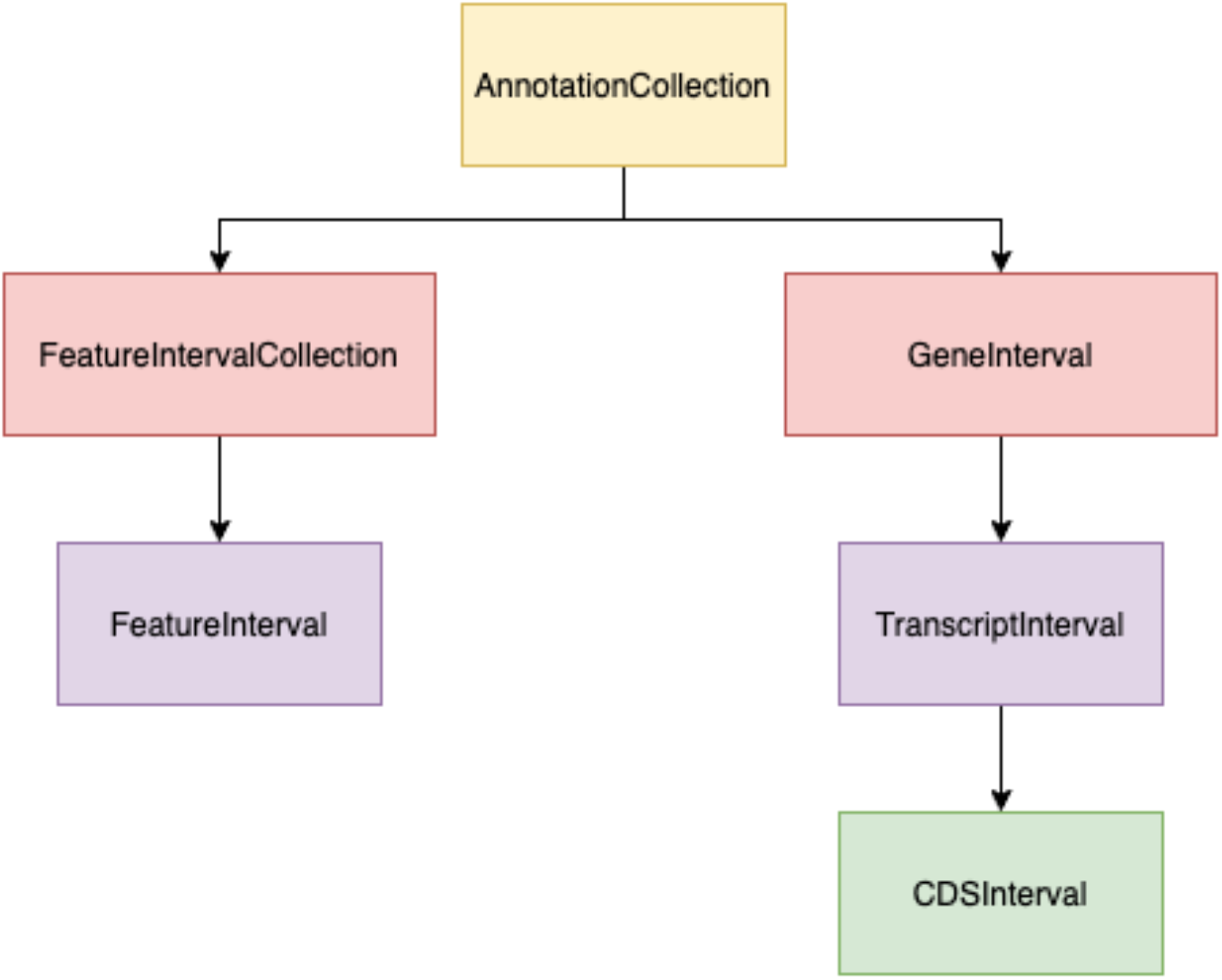
Annotation data structure. `AnnotationCollection` objects hold any number of arbitrary intervals in a contiguous genomic region on a single sequence. `AnnotationCollection` objects contain one or more `FeatureIntervalCollection` and `GeneInterval` children. `FeatureIntervalCollection` are thought of as generic regions of the genome, such as promoters or transcription factor binding sites. Both `GeneInterval` and `FeatureIntervalCollection` have one or more `TranscriptInterval` or `FeatureInterval` children respectively. `TranscriptInterval` objects have an optional child `CDSInterval` object that model their coding potential.

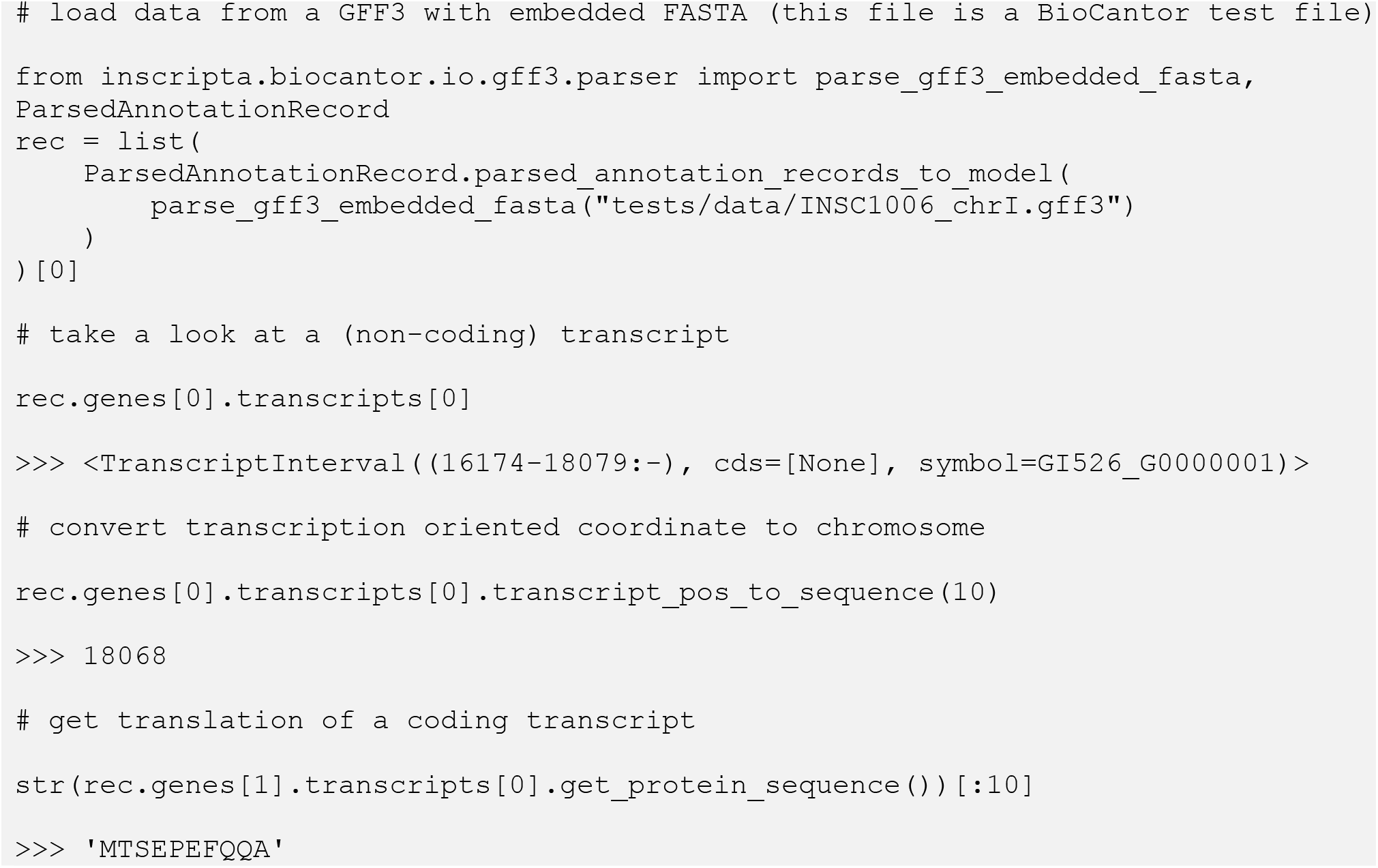
Example: Constructing and using AnnotationCollection objects.

### Data models and file formats

All of the BioCantor data structures are representable in JSON format, allowing them to be serialized to disk **(Figure 2)**. In order to facilitate building these representations, BioCantor includes parsers for GenBank and GFF3(+FASTA) **(Table 2)** format annotation files. In order to provide interoperability with common bioinformatics workflows, BioCantor data models can also be exported to GFF3, GenBank, BED, and NCBI TBL format.

**Figure 2.**
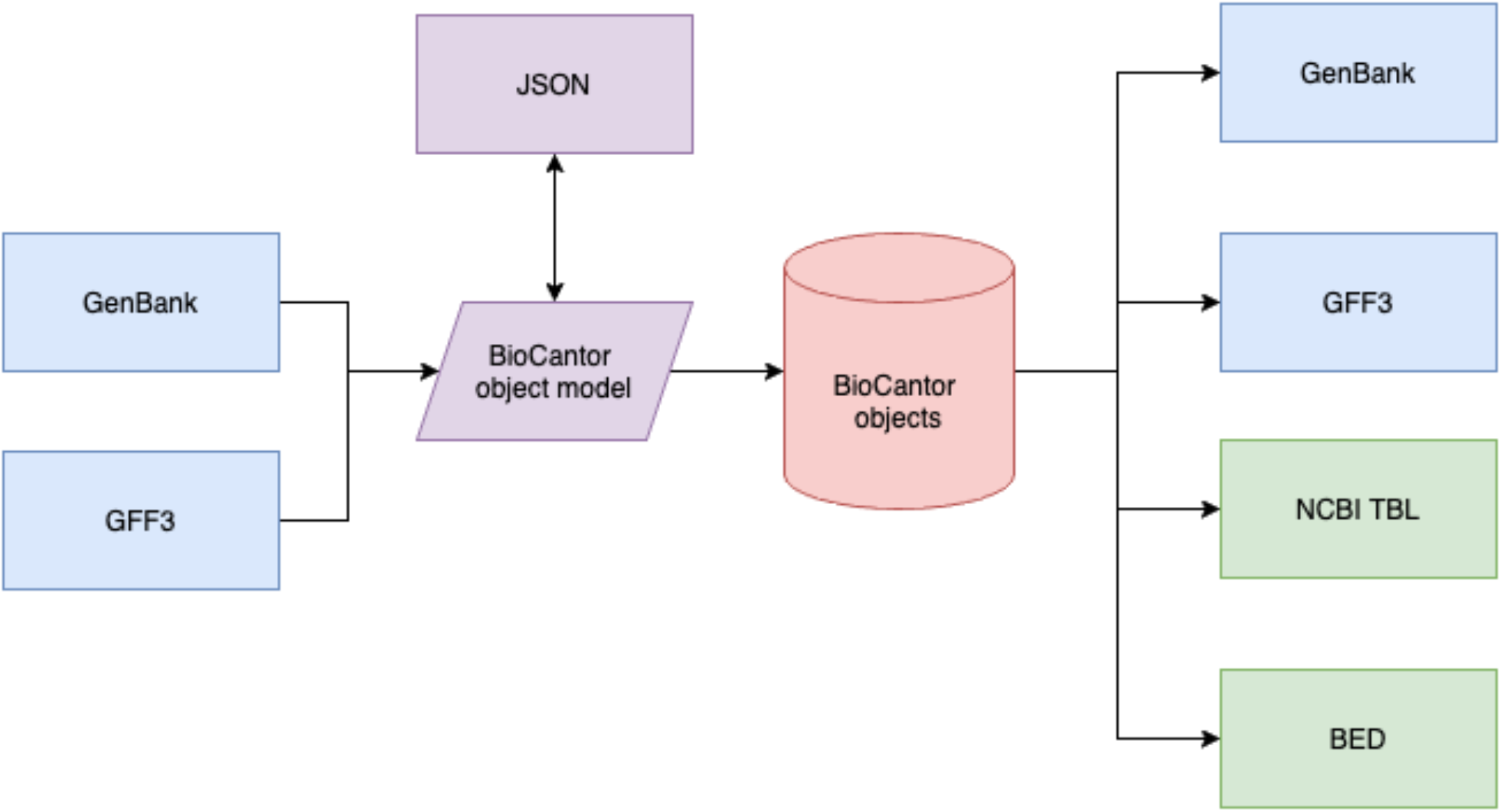
BioCantor file parsing. Parsers for GenBank and GFF3 files produce a JSON-serializable object model representation. The object model can be converted to BioCantor interval objects, with optional sequence information. These interval objects can be exported to GenBank, GFF3, NCBI Feature table (.tbl), and BED format.

**Table 2.** Annotation file parser support. BioCantor supports parsing GenBank files as well as GFF3 files with or without FASTA files. Parsing GFF3 without FASTA will produce data structures that can perform coordinate arithmetic and be exported to other file formats but lack sequence information.

| Format | Underlying Parser | Notes |
| --- | --- | --- |
| GenBank | BioPython<sup>6</sup> | Automatically associates sequence information. |
| GFF3 | gffutils<sup>7</sup> |  |
| GFF3 + separate FASTA | gffutils + BioPython | Automatically associates sequence information. |
| GFF3 + embedded FASTA | gffutils + BioPython | Automatically associates sequence information. |

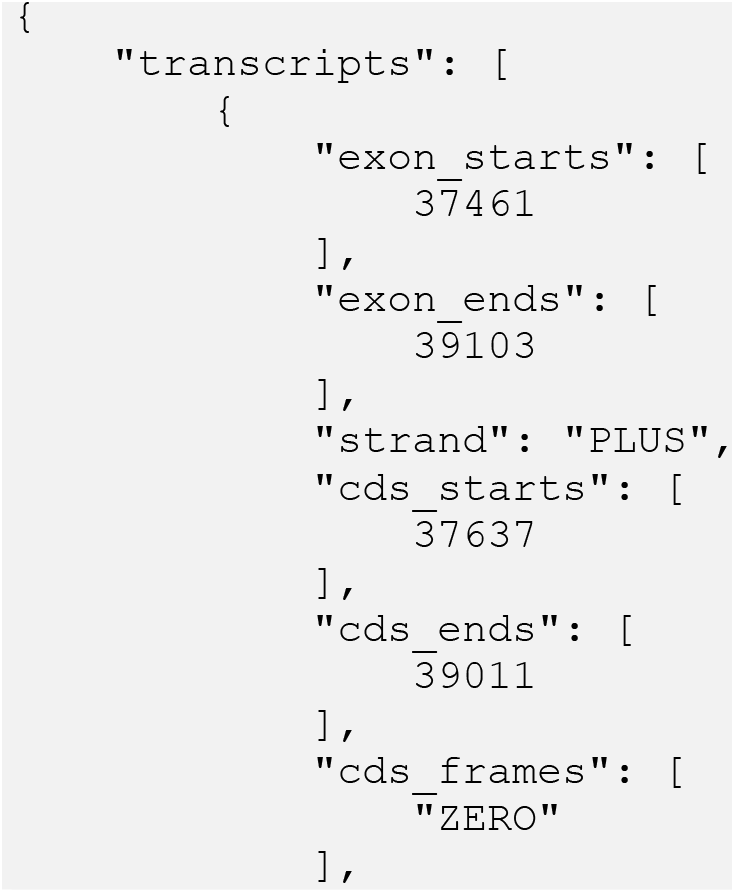
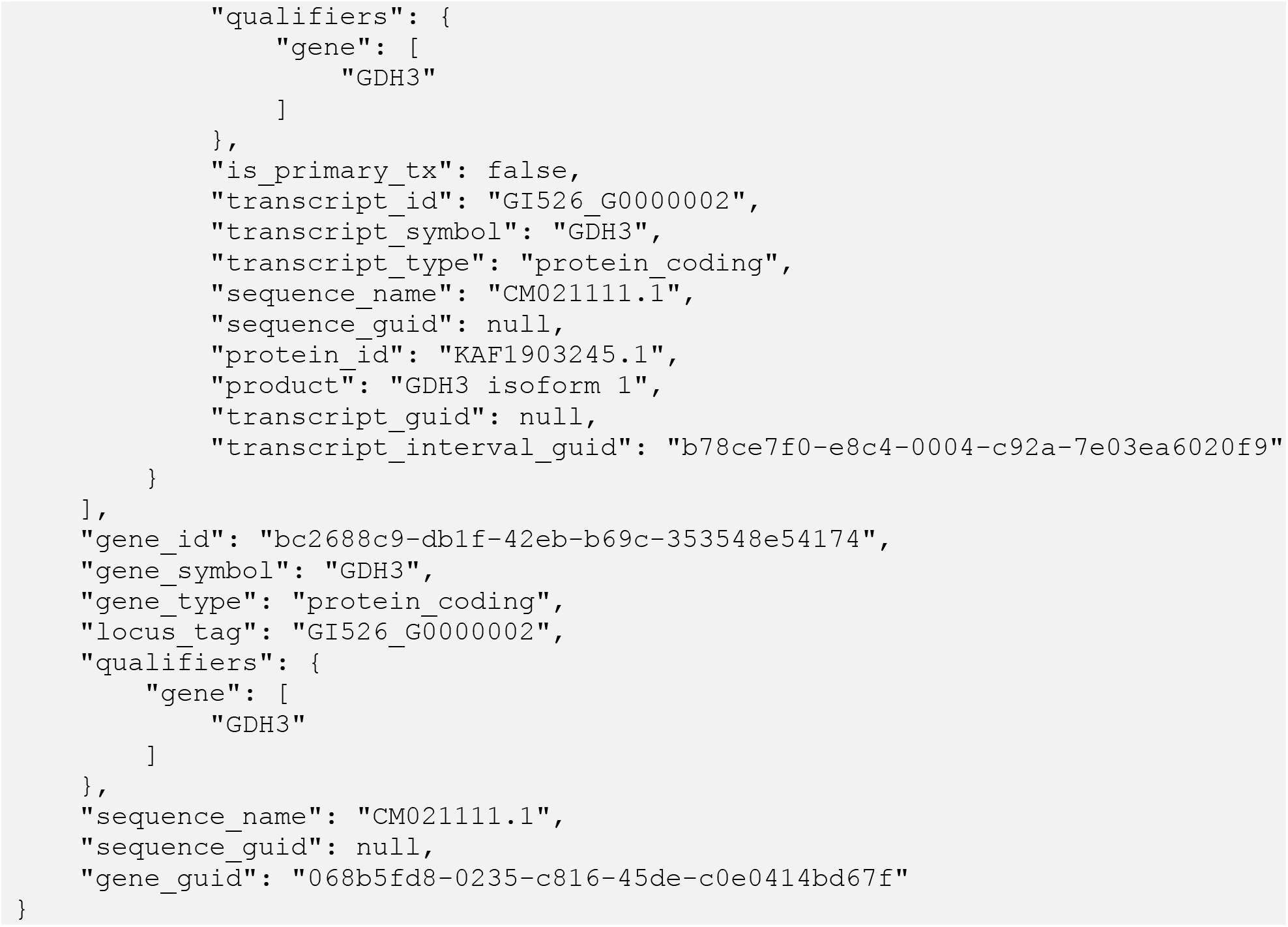
Example: JSON representation of BioCantor gene model.

## Conclusions

BioCantor enables elegant genomic workflows to be expressed in custom Python code through full end-to-end support of rich feature operations. Furthermore, the abstraction of coordinate operations lets programmers operate in multiple simultaneous coordinate systems without the need to keep track of coordinate arithmetic. The flexible paradigm can be deployed to address any use case requiring genomic feature arithmetic.

